# Machine learning prediction and experimental validation of antigenic drift in H3 influenza A viruses in swine

**DOI:** 10.1101/2020.08.07.238279

**Authors:** Michael A. Zeller, Phillip C. Gauger, Zebulun W. Arendsee, Carine K. Souza, Amy L. Vincent, Tavis K. Anderson

## Abstract

The antigenic diversity of influenza A virus (IAV) circulating in swine challenges the development of effective vaccines, increasing zoonotic threat and pandemic potential. High throughput sequencing technologies are able to quantify IAV genetic diversity, but there are no accurate approaches to adequately describe antigenic phenotypes. This study evaluated an ensemble of non-linear regression models to estimate virus phenotype from genotype. Regression models were trained with a phenotypic dataset of pairwise hemagglutination inhibition (HI) assays, using genetic sequence identity and pairwise amino acid mutations as predictor features. The model identified amino acid identity, ranked the relative importance of mutations in the hemagglutinin (HA) protein, and demonstrated good prediction accuracy. Four previously untested IAV strains were selected to experimentally validate model predictions by HI assays. Error between predicted and measured distances of uncharacterized strains were 0.34, 0.70, 2.19, and 0.17 antigenic units. These empirically trained regression models can be used to estimate antigenic distances between different strains of IAV in swine using sequence data. By ranking the importance of mutations in the HA, we provide criteria for identifying antigenically advanced IAV strains that may not be controlled by existing vaccines and can inform strain updates to vaccines to better control this pathogen.

## INTRODUCTION

Influenza A virus (IAV) is a primary respiratory pathogen in commercial swine in the United States (1). Preventing infection and transmission of the virus has proven difficult due to rapid mutation that allows the virus to evade host immune defenses and impacts the efficacy of vaccination programs by antigenic drift (2). The best approach for effective IAV control has been the development of vaccines that reflect the antigenic diversity of circulating swine IAV strains (3). This is dependent on robust sampling and sequencing of contemporary strains, which is currently achieved primarily through passive surveillance, where clinically sick pigs are sampled, and the hemagglutinin (HA) gene is sequenced and compared to vaccine antigens based on either genetic clade or sequence identity. Vaccines that include a well-matched HA can induce the production of antibodies that may provide sterilizing immunity, help reduce clinical signs, or reduce transmission (4,5). Conversely, mismatched vaccine antigens can result in vaccine failure or potentially cause enhanced disease, emphasizing the importance of careful vaccine strain selection (6).

In the United States, swine IAV is monitored by the United States Department of Agriculture (USDA) in collaboration with regional veterinary diagnostic laboratories in the National Animal Health Laboratory Network (7). These data are primarily synthesized using phylogenetic analysis (7,8), but there is no coordinated effort to characterize the phenotypic differences between circulating viruses (9). This contrasts the approach for human IAV, where vaccine antigens are selected through comprehensive genetic and antigenic characterization of seasonally circulating IAV (10). Thus, the majority of vaccine antigens in use for IAV in swine are selected based solely on the genetic clade or percent amino acid identity. This effort is fraught with risk as there are at least 16 distinct HA genetic clades of IAV in swine derived from multiple human-to-swine interspecies transmission events and subsequent evolution in the swine host (8,11). Further, there is evidence for regional patterns in HA clade persistence (8,12), and the demonstration that as few as six amino acid mutations within the HA may affect the antigenic phenotype of a virus (13,14). Consequently, there is a critical need to not only sequence and genetically characterize swine IAV, but determine what of the genetic diversity is meaningful for antigenic drift.

The antigenic properties of IAV are a manifestation of the structural interaction between IAV and host antibodies (15-18). Structural changes in the HA may alter the interaction with antibodies targeting the virus, and these changes are generally correlated with the number of accumulated amino acid mutations in the HA protein (19). Empirical data has also shown that certain amino acid mutations have a disproportionate effect on antigenic change based on the location of the amino acid in the protein structure (13,15). Though there are relatively few antigenically characterized swine IAV HA genes (9,13), this empirical data may be used to establish antigenic distances between multiple IAV in swine, and be used to gain insight on the contribution of site-specific amino acid mutations. These data can subsequently be used to assign a level of importance to specific amino acid mutations and be used to predict antigenic drift and the biological relevance of genetic diversity collected during surveillance programs.

In this study, machine learning methods were used to model the antigenic properties of IAV in swine and predict the antigenic distance between different strains using HA sequences. Modelling methods, such as the ones we present, are able to overcome the prohibitive costs and logistical challenges associated with large scale phenotypic characterization. These data can be used in combination with in-field surveillance platforms (20) as an approach for the early detection of antigenic variants and novel viruses. Additionally, these algorithms can be disseminated to swine practitioners in analytical pipelines (11,20,21) to facilitate the rational design of vaccines that include antigens that will likely protect against the circulating IAV strains. Understanding how genetic diversity, and which amino acids within the HA gene are the most important, can allow for the simulation of the antigenic evolution of swine IAV and make predictions about the persistence and circulation of future IAV strains.

## MATERIAL AND METHODS

### The swine IAV H3 antigenic reference dataset

The antigenic properties of two influenza viruses can be quantitatively compared using a hemagglutination inhibition (HI) assay. The assay is based on the ability of the hemagglutinin to agglutinate red blood cells, which express sialic acid on their cell surface (22,23). The HI antibodies raised against a homologous IAV can block the agglutination of red blood cells, even at low concentrations. Genetically different viruses often need a higher concentration of HI antibodies to prevent agglutination compared to the homologous titer. Comparing the antigenic distance between two viruses is calculated by distance *D*_*ij*_ = log_2_(*H*_*ij*_) – log_2_(*H*_*ij*_), representing a two-fold loss in HI antibody cross-reactivity between the homologous and heterologous HI antibody titers (24). These data have traditionally been used to generate pairwise antigenic distances between IAV in swine that is then visualized using multidimensional scaling to form an antigenic map (9,25,26).

The HI titers were collected from prior swine H3 HA virus characterization studies that used HI assays (23,27,28). The HI titers from new IAV selected as reference strains were collected to expand the dataset using methods described in prior literature, totaling 128 reference antigens tested against 47 reference antisera in various combinations from combined experiments (22). Distances between available HI titers were calculated by subtracting the log2 of the heterologous titer from the log2 of the homologous titer (24). Distances corresponding to the same antigen-antiserum pair were calculated as the log2 of the geometric mean as 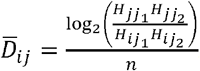.

### Training and validation of machine learning regression models

Full length HA amino acid sequences for each antigen represented in the dataset were aligned using MAFFT v7.311 (29) and then trimmed to the HA1 domain (amino acids 1-328 using the H3 HA numbering with the signal peptide removed) for subsequent analyses. Percent amino acid difference (100% - amino acid identity) was calculated between each HA pair for all combinations of sequences. Specific amino acid substitutions were not weighted to minimize model assumptions, and prior research in human IAV has suggested that these approaches may add noise to analysis (30,31). All observed site-specific amino acid substitutions in the reference data were identified and treated as bi-directional.

The regression model data was constructed with antigenic distance calculated from HI titer as the training value, with percent amino acid difference as a continuous predictor feature, and site-specific mutations as binary predictor features. Three different machine learning regression models were trained using scikit-learn (32): random forest; adaBoost decision tree; and multilayer perceptron. For each regression model, hyperparameters were tuned using a random search optimization (Supplemental Table 1). A fourth regression model was created by averaging the three prior machine learning model predictors and referred to as the ensemble model.

**Table 1.** Performance indicators for the random forest, adaBoost decision tree, multilayer perceptron, and ensemble regression models with tuned hyperparameters. Pearson correlation and root mean squared error were determined using an 80/20% split between training and test antigen data. A 10-fold cross validation based on the root mean squared error was applied.

| Performance Indicator | Random Forest | AdaBoost Decision Tree | Multilayer Perceptron | Ensemble |
| --- | --- | --- | --- | --- |
| Pearson Correlation | 0.78 | 0.77 | 0.78 | 0.80 |
| RMSE | 1.60 | 1.28 | 1.32 | 1.21 |
| 10-Fold CV (RMSE) | 1.56 ($\pm 0.29$) | 1.59 ($\pm 0.33$) | 1.76 ($\pm 0.39$) | 1.58 ($\pm 0.27$) |

Data was split into 80% training data and 20% testing data groups to calculate the Pearson correlation and root mean squared error. Additionally, 10-fold cross validation was used to assess the root mean squared error (Table 1). Given the sparsity of antigenic data available, a leave-one-out cross validation approach was employed to generate a distribution of prediction error for each model (Figure 1). Each antigen included in the training set (n = 128) was iteratively excluded from the training set and distances were predicted using each of the four regression models. The error was calculated as the absolute value of difference between the predicted distance and the empirical distance.

**Figure 1.**
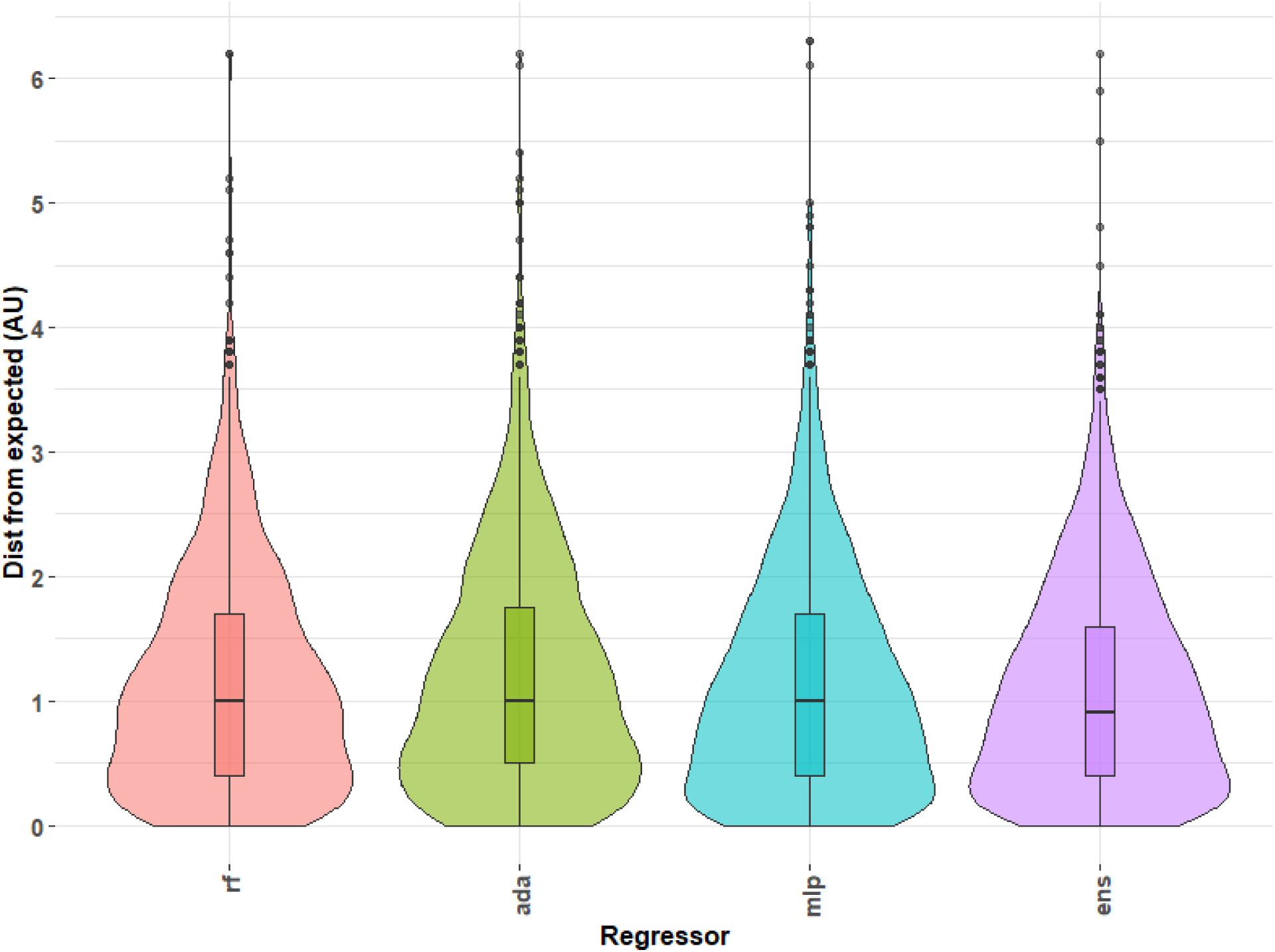
Distribution of error calculated for the predicted antigenic distance compared to actual antigenic distance as predicted by machine learning models and hemagglutination inhibition assays, respectively. Three regression models were used to predict distances from empirically determined antigens using hemagglutination inhibition titers in a leave-one-out approach: random forest regression (rf), adaBoost decision tree regression (ada), and multilayer perceptron (mlp) regression. All three predictions were combined into an ensemble (ens) to prevent overfitting and to minimize errant predictions by averaging across predictions from all models. Approximately 25% of the data has 0.5 antigenic units (AU) of error or less, 50% of the data has 1 AU of error or less, 75% of the data being less than 2 AU of error. Maximum error for outliers exceeded 6 AU.

### Mapping antigenic predictions onto phylogenetic trees

Maximum-likelihood phylogenetic trees were created to assess antigenic distance predictions of genetically similar sequences of the test antigen sequence compared to the reference sequence. Sequences were aligned using MAFFT v7.311 (29) and phylogenetic trees were inferred using FastTree v2.1.10 (33). Trees were annotated using FigTree v1.4.3 (34) with each tree rooted to a reference strain and sorted in ascending order relative to inferred evolutionary relationship. Each tip within the tree was color-coded based on the antigenic motif designated by H3 numbering positions 145, 155, 156, 158, 159, and 189 as prior work identified these sites as significant for antigenic phenotype (15). Branches were annotated with the ensemble-predicted antigenic distance relative to the root. Trees were pruned to 30 leaves to facilitate viewing.

### Determining the relative importance of genetic mutations

Random forest regression models provide a natural ranking system of feature importance (35). The importance of each predictor feature was calculated by the decrease in the node variance after fitting the random forest model. The feature rankings for the random forest regression model were analyzed to assess the biological importance of observed mutations in the swine H3 antigenic reference dataset. The significance of each amino acid position in the HA was determined by summing the mutation-based features grouped by the position they represented. The resultant significance of each amino acid was projected onto a protein model of a human H3 HA gene A/Victoria/361/2011 obtained from the Research Collaboratory for Structural Bioinformatics (4O5N) (36).

### Empirical validation of machine learning regression models

The H3 HA amino acid sequences of uncharacterized IAV in swine submitted to NCBI GenBank from the Iowa State University Veterinary Diagnostic Lab from January 2016 to August 2018 were collected and clustered by phylogenetic clade (7,11). The HA gene sequences were trimmed to the HA1 domain (positions 1-328 using H3 numbering with the signal peptide removed). The HA1 sequences were compared against all antigenically characterized sequences to calculate percent amino acid difference and compare the presence or absence of site-specific amino acid mutations. Site-specific amino acid mutations absent from the training set were not considered in additional analyses. The antigenic distance from each uncharacterized HA gene to each reference antigen was predicted using the previously described four trained regression models.

A selection of four contemporary IAV were selected as test antigens to be antigenically characterized with in vitro HI assays to validate the regression models using their HA genes. We selected these HA genes from within the H3-Cluster IVA genetic clade, as: a) this is a significant genetic clade that is frequently detected in diagnostic submissions to the Iowa State University Veterinary Diagnostic Lab (11); b) this genetic clade was responsible for more than 300 zoonotic infections from 2012 to present; c) there was a significant amount of uncharacterized data within the last 2 years (n = 299 from 2018 to present, representing 8% of sequenced HA genes). Since the ensemble predictions demonstrated the least error in the analyses above, antigenic distances of 106 H3-cluster IVA viruses were predicted against a panel of 44 available antisera using this model. We selected four test antigens/antisera prediction pairs within this genetic clade based on the following criteria: near amino acid sequence identity (≥ 98%) and near predicted ensemble antigenic distance measured in antigenic units (AU) (≤ 2AU); a near identity and far antigenic distance (≥ 3AU); far identity (≤ 95%, ≥ 90%) and near antigenic distance (≤ 2AU); or far identity (≤ 95%, ≥ 90%) and far antigenic distance (≥ 3AU) (Figure 2, Table 3).

**Table 3.**
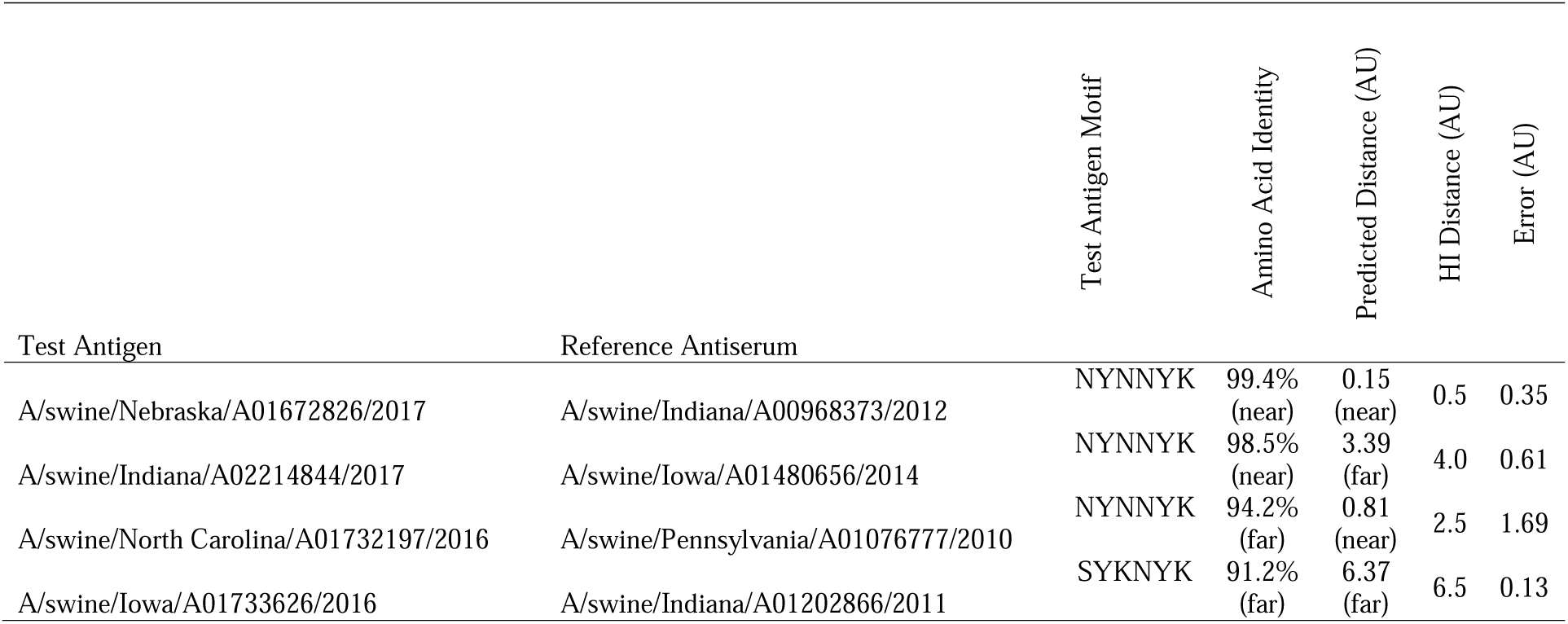
Predicted and measured antigenic distances between test antigens and reference strain antisera using the model to calculate the predicted distance and hemagglutination inhibition (HI) titers to calculate the empirical distance in antigenic units. Error is calculated by taking the absolute value of the predicted distance subtracted from the empirical distance.

| Test Antigen | Reference Antiserum | Test Antigen Motif | Amino Acid Identity | Predicted Distance (AU) | HI Distance (AU) | Error (AU) |
| --- | --- | --- | --- | --- | --- | --- |
| A/swine/Nebraska/A01672826/2017 | A/swine/Indiana/A00968373/2012 | NYNNYK | 99.4%<br>(near) | 0.15<br>(near) | 0.5 | 0.35 |
| A/swine/Indiana/A02214844/2017 | A/swine/Iowa/A01480656/2014 | NYNNYK | 98.5%<br>(near) | 3.39<br>(far) | 4.0 | 0.61 |
| A/swine/North Carolina/A01732197/2016 | A/swine/Pennsylvania/A01076777/2010 | NYNNYK | 94.2%<br>(far) | 0.81<br>(near) | 2.5 | 1.69 |
| A/swine/Iowa/A01733626/2016 | A/swine/Indiana/A01202866/2011 | SYKNYK | 91.2%<br>(far) | 6.37<br>(far) | 6.5 | 0.13 |

**Figure 2.**
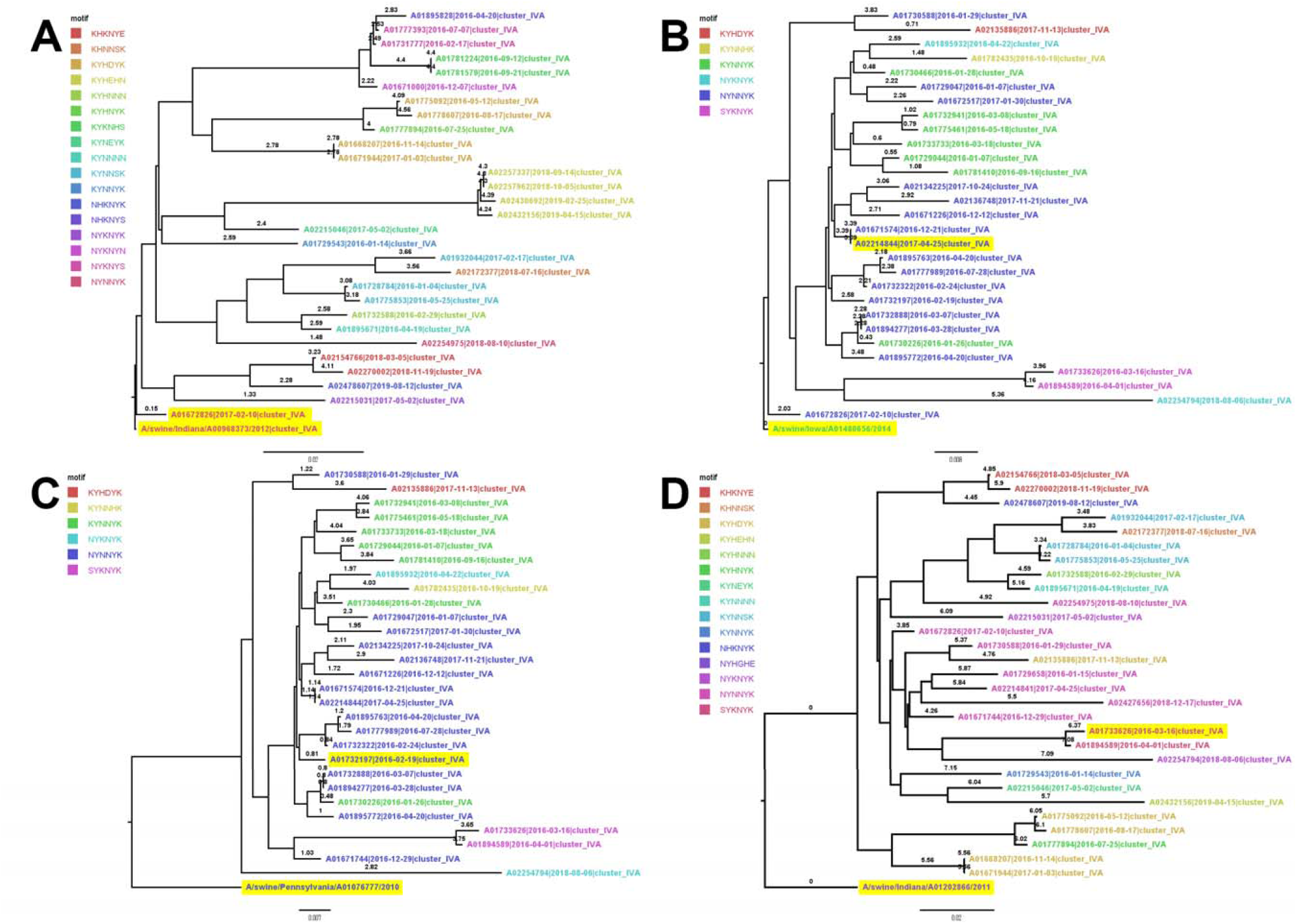
Phylogenetic trees of test antigens rooted to their reference strain. A) Phylogenetic tree of test antigen A/swine/Nebraska/A01672826/2017 and reference strain A/swine/Indiana/A00968373/2012, representing a near predicted antigenic distance prediction (0.16 AU) for two strains of near amino acid identity (99.4%). B) Phylogenetic tree of test antigen A/swine/Indiana/A02214844/2017 and reference strain A/swine/Iowa/A01480656/2014, representing a far predicted antigenic distance prediction (3.3) for two strains of near amino acid identity (98.5%). C) Phylogenetic tree of test antigen A/swine/North Carolina/A01732197/2016 and reference strain A/swine/Pennsylvania/A01076777/2010, representing a near predicted antigenic distance prediction (0.31) for two strains of far amino acid identity (94.2%). D) Phylogenetic tree of test antigen A/swine/Iowa/A01733626/2016 and reference strain A/swine/Indiana/A01202866/2011, representing a far predicted antigenic distance prediction (6.33) for two strains of far amino acid identity (91.2%). Branches of the phylogenetic tree were annotated with the predicted antigenic distance from the ensemble regression model (both test antigen and reference strain are highlighted). Each tree is pruned to 30 sequences. Influenza strains are colored by the antigenic motif formed by amino acid positions 145, 155, 156, 158, 159, and 189: these positions, located near the ligand binding site of the hemagglutinin protein, have been noted to affect the antigenic interactions of the protein.

The four selected antigen/antisera pairs were tested via HI assay. HI assays were conducted as previously described (23) with empirical distances calculated by taking the log2 of the heterologous titer subtracting from the log2 of the homologous titer. Empirical distances were compared against predicted values by subtraction.

## RESULTS

### Machine learning model performance

Comparison of the empirical antigenic distances against the predicted values indicated that the Pearson correlation for all regression models was within a range between 77%-80% (Table 1). The root mean squared error (RMSE) was between 1.21 – 1.60 antigenic units of error depending on the model. Ten-fold cross validation of the random forest, adaBoost decision tree, and multilayer perceptron regression models had an RMSE of 1.56 ± 0.29, 1.59 ± 0.33, and 1.76 ± 0.39 respectively. The leave-one-out cross validation demonstrated that for all models, 25% had ≤ 0.5 AU, 50% had ≤ 1.0 AU, and 75% had ≤ 1.7 AU distance error. The maximum observed error was 6.3 AU, with each model producing errors > 6.0 AU (Figure 1).

### Mapping antigenic predictions onto phylogenetic trees

Four trees were built with sequences genetically similar to each test antigen (Figure 2). Trees were annotated with an amino acid motif based on positions 145, 155, 156, 158, 159, and 189 as these sites have been found to have a disproportionate effect on the observed antigenic phenotype in both human and swine H3 (14). The antigenic motif between test antigen A/swine/Nebraska/A01672826/2017 and reference antiserum A/swine/Indiana/A00968373/2012 match, both being NYNNYK (Figure 2A). The antigenic motif of test antigen A/swine/Indiana/A02214844/2017 was NYNNYK, while reference antiserum A/swine/Iowa/A01480656/2014’s motif was KYNNYK, differing at position 145 (Figure 2B). The antigenic motif between test antigen A/swine/North Carolina/A01732197/2016 and reference antiserum A/swine/Pennsylvania/A01076777/2010 match, both being NYNNYK (Figure 2C). The antigenic motif of test antigen A/swine/Iowa/A01733626/2016 was SYKNYK, while reference antiserum A/swine/Indiana/A01202866/2011’s motif was NYHGHE, differing at positions 145, 156, 158, 159, 189 (Figure 2D).

### Empirical validation of the predicted antigenic distance predictions

The predicted ensemble distances of the selected test antigens were validated via HI assay (Supplemental Table 2). Test antigen A/swine/Nebraska/A01672826/2017 was predicted to be 0.15 AU from reference strain A/swine/Indiana/A00968373/2012, sharing 99.4% amino acid identity between the HA1 segments of the HA (Table 2). Both the reference and test antigens were from the H3-cluster IVA clade (Figure 2A), and this pairing represented the near identity and near antigenic distance prediction. The amino acid differences between the reference strain and the test antigen were at M10T and R208I (Table 2). The HI assay demonstrated the antigenic distance between the reference strain antiserum and test antigen was 0.5 AU (Table 3) with an error between the predicted distance and the empirical distance of 0.35 AU.

**Table 2.** Amino acid mutations detected between test antigen and reference strains used for the model validation.

| Test Antigen | Reference Strain | Amino Acid Changes |
| --- | --- | --- |
| A/swine/Nebraska/A01672826/2017 | A/swine/Indiana/A00968373/2012 | M10T, R208I |
| A/swine/Indiana/A02214844/2017 | A/swine/Iowa/A01480656/2014 | G49S, E83K, V112I, K145N, S289P |
| A/swine/North Carolina/A01732197/2016 | A/swine/Pennsylvania/A01076777/2010 | T10M, E83K, V106S, A107T*, V112I, T117N, N124S, K142S, A163E, M168V, N173K, I196V, T203I, P273H, G275D, N276E, K278N, R299K, V304A |
| A/swine/Iowa/A01733626/2016 | A/swine/Indiana/A01202866/2011 | I29L, G50R, E83K, S107T, T117N, S124N, A131D, D133G, R137N, S138T, R140K, G144V, N145S, H156K, G158N, H159Y, A163E, L164Q, T167A, N173K, E189K, S193N, V196A, I203V, R220V, R269K, S273H, N276E, R299K |

\* Changes not accounted for by regression models

Test antigen A/swine/Indiana/A02214844/2017 was predicted at 3.39 AU from reference strain A/swine/Iowa/A01480656/2014, sharing 98.5% amino acid identity between the HA1 segments. Both the reference strain and test antigens are from the H3-cluster IVA clade (Figure 2B), and this pairing represents near identity but far antigenic distance prediction. There were 5 amino acid differences between the reference strain and test antigen (Table 2). The HI assay found a distance of 4.0 antigenic units between the test antigen and reference antiserum and an error of 0.61 AU between empirical and predicted distances.

Test antigen A/swine/North Carolina/A01732197/2016 was predicted at 0.81 AU from reference strain A/swine/Pennsylvania/A01076777/2010, sharing 94.2% amino acid identity between the HA1 segments. The test antigen was selected from the H3-cluster IVA clade and the reference strain from the H3-cluster IV clade (Figure 2C), and this pairing represented a distant identity that was predicted to be antigenically similar. There were 19 amino acid differences between the reference strain and test antigen, with the A107T mutation being the only position not accounted for in the trained model (Table 2). The HI assay demonstrated an average antigenic distance between reference antiserum and test antigen of 2.5 AU, with a prediction error of 1.69 AU.

A/swine/Iowa/A01733626/2016 was predicted at 6.37 AU from reference strain A/swine/Indiana/A01202866/2011, sharing 91.2% amino acid identity between the HA1 segments. The test antigen is from the H3-cluster IVA clade of virus and reference strain from the H3-cluster IVC clade (Figure 2D). This pairing represents a far identity and far predicted antigenic distance prediction. There were 29 amino acid differences between the reference strain and test strain (Table 2). The HI assay demonstrated 6.5 antigenic units between test antigen and reference antiserum, giving an error of 0.13 AU between empirical and predicted distances.

### Ranking of predictor features

Random forest regression ranks user-selected features by a metric of importance, calculated by the decrease in the node variance and normalized across the forest for a single model run (Supplemental Table 3). The highest-ranking features were stable across runs as they had a consistent decrease in their average variance, though these metrics were susceptible to starting conditions (data provided at https://github.com/flu-crew/antigenic-prediction). The most important feature in predicting the antigenic distance between two strains was amino acid identity within the HA1, accounting for 31.4% of the importance. Transitions between K and N at position 145 accounted for 8.1% of the model importance and was ranked as the most important amino acid mutation. However, transitions between K and S and N and S at the same position 145 received lower ranking in model importance (totaling 0.2% importance cumulatively), demonstrating that the context of the positional mutation is important. Features I202V and R222W (representing bi-directional mutations) ranked at 5.4% and 5.2% importance respectively. The remainder of the features in the models accounted for less than 3% of the model on an individual basis (Figure 3, Supplemental Table 3), with the next ten bidirectional mutations in order of importance as H75Q, R137Y, D101Y, E62K, I25L, P289S, D133N, E189K, K92T, and H159Y (Figure 3). Projecting the cumulative importance of each amino acid position on an H3 crystal structure indicated that position 145, the most important position in the model, is located in the groove of the active site (Figure 4). Other sites of higher importance were more likely to be observed on the solvent facing side of the trimer. Amino acid position 202 was an exception as it was ranked as of high importance but was located on the inside of the trimer.

**Figure 3.**
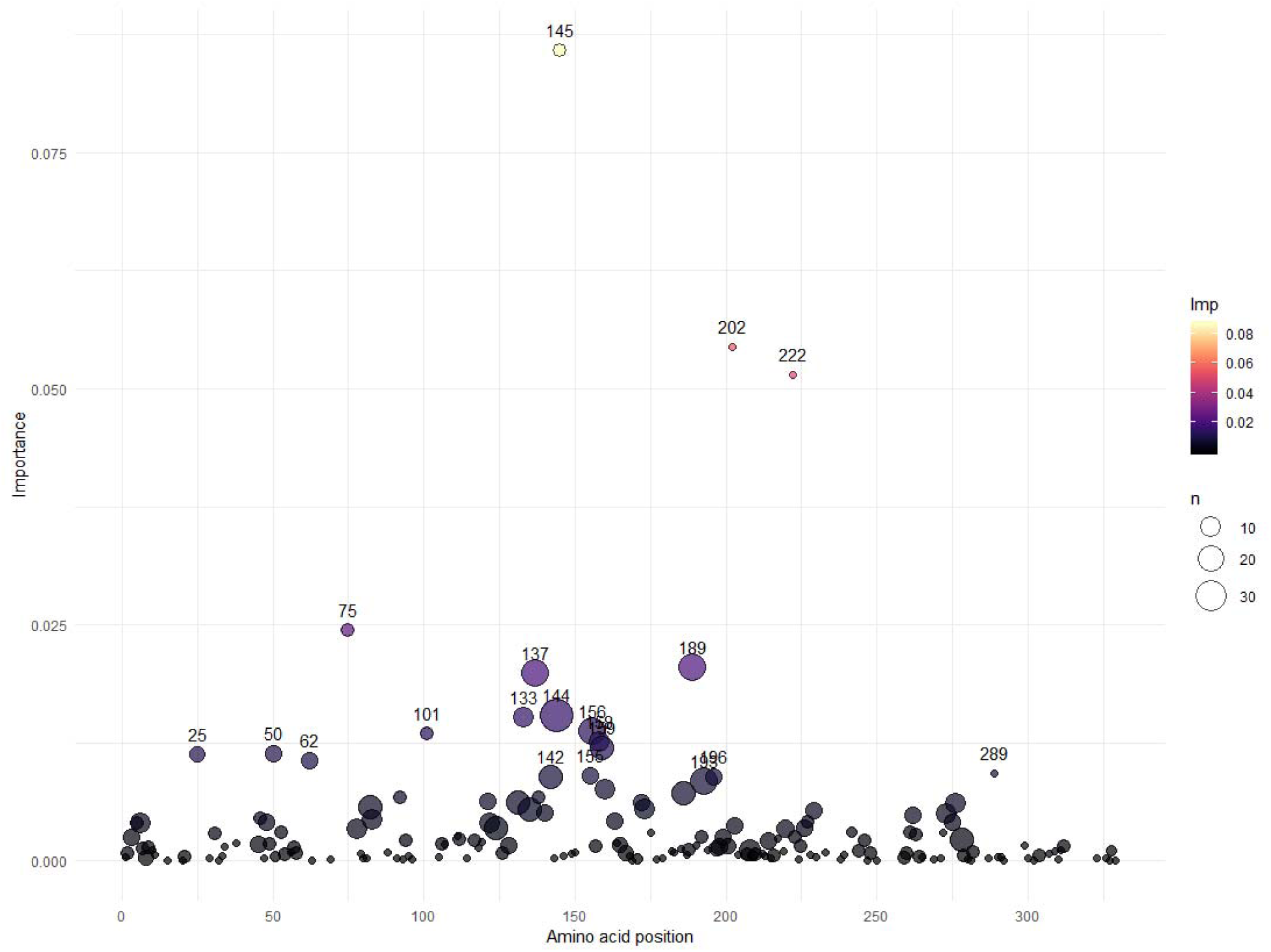
Rank of amino acid location importance by the cumulative summation of importance per site mutation as determined by random forest regression. Amino acid position using H3 numbering is reported on the x-axis. The importance for each site-specific mutation is summed per site and displayed on the y-axis using a color scale. The size of the circle is relative to the number of mutations observed in the training set per site. Identity was the highest-ranking feature, with an importance of 0.312, but is not displayed on the graph. The top ten amino acid transition features in order of importance are K145N, R222W, I202V, H75Q, I25L, R137Y, D101Y, E62K, P289S, and D133N. The top ten amino acid sites in order of cumulative importance are 145, 222, 202, 75, 189, 137, 25, 133, 144, and 156.

**Figure 4.**
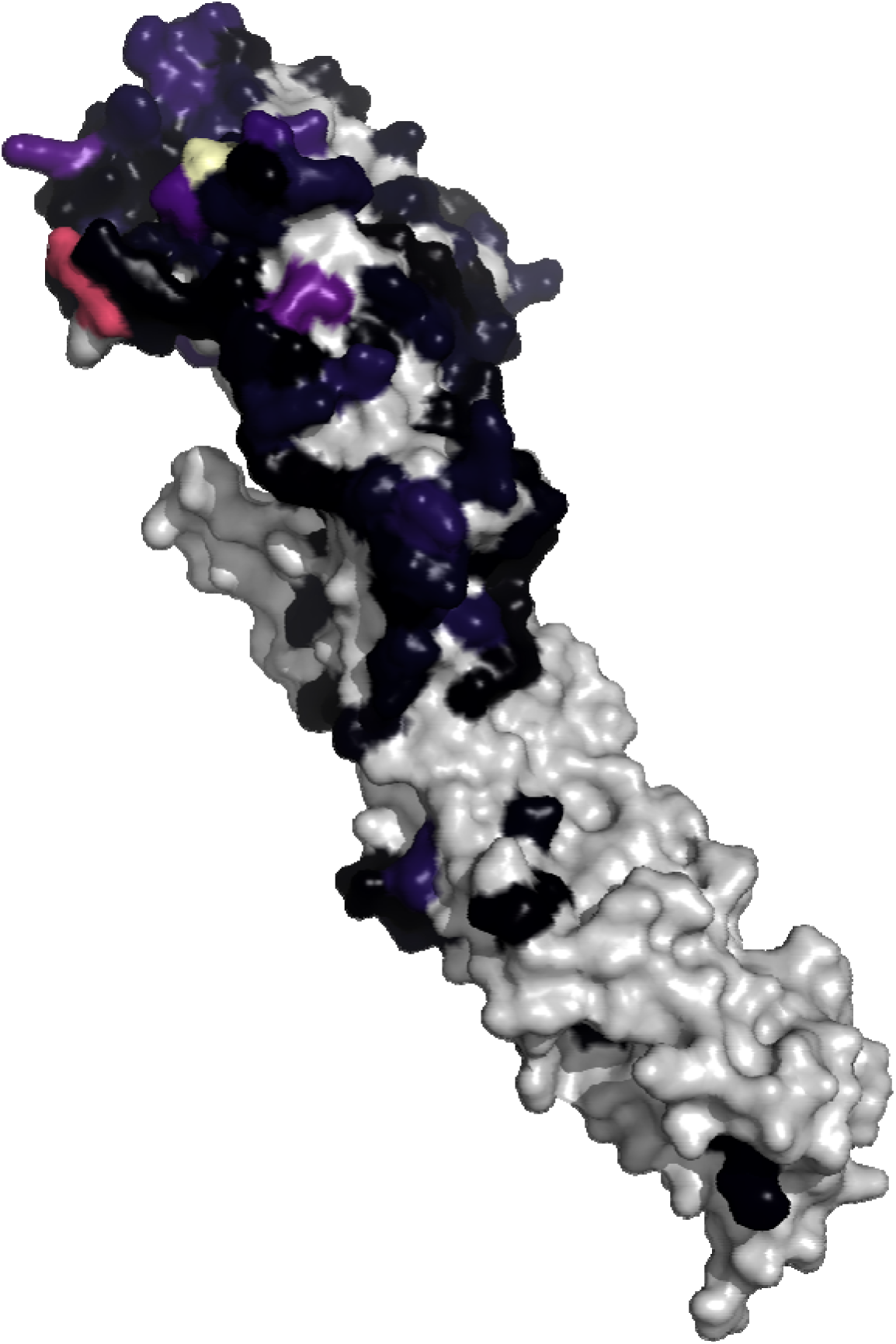
Projection of feature importance on a monomer of the A/Victoria/361/2011 hemagglutinin (HA) protein (RCSB 4O5N). The significance of each amino acid position in the HA was determined by summing the substitution-based features grouped by the position they represented. Significant positions were projected onto a hemagglutinin protein model of the human H3. The importance for each site-specific mutation is summed per site and projected onto the hemagglutinin protein model of the human H3. Higher color intensity represents a larger calculated importance. Positions with no data were colored gray.

Of the 728 features included in the model, amino acid identity and the sum of the top ten amino acid mutation features of the model accounted for 58.3% of the importance. Identity and the top 253 amino acid mutation features accounted for 95% of the calculated importance, whereas the top 397 features accounted for 99% of the calculated importance.

## DISCUSSION

In this study, a model was developed to computationally estimate antigenic distances between different IAV in swine based on amino acid sequence using non-linear machine learning methods. The method leverages data that was generated from previous antigenically characterized IAV strains in swine to train regression models. After in silico validation, the models were used to predict the antigenic distance between paired IAV strains based on their amino acid identity and the mutations present between each strain. Finally, the antigenic distance predictions were experimentally confirmed by comparing the distance between homologous and heterologous hemagglutination inhibition (HI) titers. Predicting antigenic distances between two genetically related but antigenically different IAV reduces the number of HI assays that are required to perform the analysis and select candidate strains for a vaccine when sufficient antigenic distance between two IAV suggests a loss in antibody cross-reactivity.

We experimentally validated our model using four test antigens, with the empirical data demonstrating predictions generally had an error less than 1 AU. The error between the test antigen and reference antiserum representing a near identity with a near predicted antigenic distance was 0.35 AU (Table 3). The distance between the same test antigen and reference antiserum HI titers was calculated at 0 and 1 AU (Supplemental Table 2), giving an average distance of 0.5. It should be noted that the HI assay is a discrete measure whereas the prediction is continuous, thus an error less than 1 AU is not biologically meaningful. Additionally, because of the discrete nature of the HI assay, the 0.5 AU error is negligible as the true antigenic distance is somewhere between 0 and 1 AU. The near identity with a far predicted antigenic distance had a wider range between the two sera’s HI titers 3 and 5, but the predicted distance 3.39 was within this range, and had an error of 0.61 AU from the average of 4 AU. The far identity with a near predicted antigenic distance had HI titers of 2 and 3, with a predicted distance of 0.81, giving an error of 1.69 AU from the average of 2.5 AU. Although the error was higher than the other predictions, the ensemble prediction was able to discern that these two strains were more antigenically similar than would be predicted based on sequence similarity alone. For the far identity and far predicted antigenic distance test antigen and reference antiserum pair, the predicted distance was 6.37 and the empirical distance was 6.5. Given the raw antigenic distances calculated from the pair of titers were 6 and 7 for the two serum samples, the real distance is likely somewhere between the two values. Consequently, our approach that was developed using a small IAV in swine empirical dataset made predictions that in the majority of cases are useful in biological applications Machine learning methods can assign importance to the position and context of amino acid mutations, allowing biological interpretation. Assessing the importance of the random forest model revealed that both the position and context of the amino acid mutation contributed to observed antigenic phenotype. While sequence difference had the highest importance in the random forest model, further assessment of the model revealed unequal weight between amino acid positions representing different mutations. An example of this dynamic was H3 HA position 145 where a mutation between K and N bidirectionally was ranked as the most important amino acid mutation feature. Other observed mutations at position 145 between K and S and N and S were ranked as less important, matching the biological nuances that have been observed with empirical testing and other computational predictions (15,43). Literature reports suggested that the conservation of biochemical properties of the amino acid mutation may also have some effect on the observed antigenic change (15,19). Unequal weighting of mutations in the model suggests antigenic distance may help improve vaccine antigen selection when compared to HA sequence comparison alone, as this approach captures not only sequence homology but how amino acid can influence antigen-antibody interactions.

Our method identified sites that had a major impact of the antigenic phenotype of swine IAV. The majority of these sites were located on the solvent exposed surface of the HA protein and in antibody epitopes that have been identified in human IAV (Figure 4) (50,51). Interestingly, the profile of positional feature importance displayed some differences to prior literature describing human H3N2 IAV. While there was considerable overlap between the positions in our model with the highest cumulative importance (Supplemental Table 3) compared to the positions in the JRFR algorithm (positions 62, 121, 131, 133, 135, 137, 142, 144, 145, 155, 156, 158, 159, 172, 173, 189, 193, 196, 276), the relative importance of these predictor features varied. Specifically, position 189 was the most important site in human H3 with ferret antisera, whereas our model identified position 145 as the most important position in swine H3 with swine sera (31). These differences of importance may be reflective of host specific interactions. Additionally, the distribution of importance was more evenly spread across the JRFR model whereas in the model presented here a small number of sites had disproportionate importance. Direct sequence comparison and sequence homology remain the standard approach to determining swine IAV vaccine control strategies; our data supports this approach but suggests that consideration of the location and context of mutation is more important than crude measures of sequence homology.

This work adds to a growing body of literature that aims to quantitatively predict antigenic phenotypes of IAV from the sequence without requiring HI titers for each IAV strain (19,31,42-44). Similar methodologies have been implemented for use with other viruses such as Dengue virus, where neutralizing titer distances have been predicted based on amino acid differences (45). To the best of our knowledge, prior approaches to calculate antigenic distances between IAV were trained and tested on human IAV strains where the HA genes are characterized by phylogenetic trees with a single thick trunk with short interspersed branches with far less cocirculating genetic diversity (46-48). Antigenic data for the human IAV strains used in prior approaches was generated using ferret antisera with the caveat that human and ferret immune systems potentially interact differently with the viral antigenic phenotype (49). Compared to IAV circulating in humans, HA gene phylogenetic trees from endemic IAV circulating in swine demonstrate multiple genetic clades within the same subtype that are derived from multiple human-to-swine spillover events across the last 100 years (7,39). The large genetic diversity of strains coevolving within the swine population has resulted in a similarly large breadth of antigenic diversity and evolution. Consequently, a broad range of HI assays including many genetically different IAV are needed to capture assess antigenic diversity of IAV circulating within swine. The scale of these studies has been difficult in the swine IAV research community, and there is a sparsity of antigenic characterization of IAV in swine frequently with large gaps of time between characterizations. This has the unfortunate consequence of potentially misrepresenting the antigenic diversity of swine IAV and can make it difficult to improve our understanding of evolution of IAV in swine (19,42,45).

The process and methodology we present has potential to help select vaccine IAV candidates when antigenic distance suggests a loss of cross-protection with current vaccine strains. Our process included a robust analysis of prediction error and was able to identify the limits of the models. Using 10-fold cross validation, our ensemble model had a higher RMSE when compared to a different machine learning approach developed for human IAV by Yao et al. (2017) (31). This approach used a Joint Random Forest Regression (JRFR) algorithm that also included substitution matrices for predicting antigenic distances and had a RMSE < 1.0 (31). A linear mixed-effects model employed by Harvey et al. (2016) (42) for human IAV, also had better performance than our model but this used different datasets and had a different application. The strength of our approach is that our predictions that in the majority of cases would be useful in biological applications. Leave-one-out cross validation demonstrated 54% of the predictions made with the ensemble model were at or below 1 AU of error, and 86% were below 2AU of error where <2AU distance is frequently used to indicate biological equivalence. Further, our ensemble of non-linear regression methods were chosen due to their robustness against collinearity. Several prior machine learning methods implement linear regression, despite the relationship between amino acid mutation being non-linear and not strictly additive (19,44). Linear models can mitigate issues of collinearity by implementing approaches such as ridge regression in antigen-bridges (43), or lasso regression used by nextstrain (19,45), but these approaches may result in models that are more difficult to interpret biologically. Our random forest approach was able to identify the top 10 features accounting for 58.3% of the antigenic phenotype (253 features were needed to account for 95% importance), generating explicit predictions on when mutation of the HA gene may result in antigenic drift and reduced vaccine efficacy.

This study implemented a non-linear machine learning approach to predict antigenic distances between IAV in swine based on HA1 sequence, and experimentally validated the model predictions. Our validation with HI assays using test antigen and reference strains demonstrated that this computational approach can be used to determine antigenic differences between IAV without requiring extensive HI testing in laboratories. It is currently impractical to antigenically characterize all strains of IAV isolated from swine, and our work shows that the antigenic phenotype can be reasonably predicted from genetic sequence. The performance of our approach was sufficient even though it was parametrized with a limited empirical dataset; it seems feasible that prediction can be improved as more empirical data is made available. Due to multiple introductions of IAV into swine from human and avian sources, the genetic diversity of IAV in swine exceeds what is observed for human IAV strains (11,39,40). The genetic diversity of IAV in swine is also confounded by transportation patterns that move regional IAV strains with swine to new geographic locations where additional antigenic drift and reassortment with endemic strains may occur (41). Consequently, this method can aid in IAV in swine vaccine design efforts, which currently do not have an integrated and comprehensive system such as the World Health Organization’s (WHO) global influenza surveillance program for IAV in humans (37). Providing accurate methods such as ours that predict antigenic distances of IAV in swine increase the ability of swine producers and veterinarians to make informed decisions regarding vaccine antigens with broad application across IAV in swine to help maintain swine herd health.

## AVAILABILITY

Data and code used in this research are available in a GitHub repository (https://github.com/flu-crew/antigenic-prediction)

## ACKNOWLEDGEMENT

We gratefully acknowledge pork producers, swine veterinarians, and laboratories for participating in the USDA Influenza A Virus in Swine Surveillance System and publicly sharing sequences in NCBI GenBank.

## FUNDING

This work was supported by the Iowa State University Presidential Interdisciplinary Research Initiative; the Iowa State University Veterinary Diagnostic Laboratory; the U.S. Department of Agriculture (USDA) Agricultural Research Service (ARS project number 5030-32000-120-00-D); an National Institute of Allergy and Infectious Diseases (NIAID) at the National Institutes of Health interagency agreement associated with the Center of Research in Influenza Pathogenesis, an NIAID funded Center of Excellence in Influenza Research and Surveillance (grant HHSN272201400008C to A.L.V.); the USDA Agricultural Research Service Research Participation Program of the Oak Ridge Institute for Science and Education (ORISE) through an interagency agreement between the U.S. Department of Energy (DOE) and USDA Agricultural Research Service (contract number DE-AC05-06OR23100 to Z.A.W. and C.K.S.); the Department of Defense, Defense Advanced Research Projects Agency, Preventing Emerging Pathogenic Threats program (contract number 022092-00001 to P.C.G.; ARS project number 5030-32000-120-30-I to T.K.A. and A.L.V.); and the SCINet project of the USDA Agricultural Research Service (ARS project number 0500-00093-001-00-D). Funding for open access charge from the U.S. Department of Agriculture (USDA) Agricultural Research Service (ARS project number 5030-32000-120-00-D). The funders had no role in study design, data collection and interpretation, or the decision to submit the work for publication. Mention of trade names or commercial products in this article is solely for the purpose of providing specific information and does not imply recommendation or endorsement by the USDA, DOE, or ORISE. USDA is an equal opportunity provider and employer.

## CONFLICT OF INTEREST

The authors report no conflicts of interest.

## Notes

https://github.com/flu-crew/antigenic-prediction

### Competing Interest Statement

The authors have declared no competing interest.

