## Supplemental Tables S1-S3 for "Machine learning prediction and experimental validation of antigenic drift in H3 influenza A viruses in swine"

Table S1. Hyperparameter tuning for each regression model. Random search was used to optimize select parameters of the random forest, adaBoost decision tree, and multilayer perceptron models using 5-fold cross validation. The high and low values of the search space are provided as well as parameter values from the best predictor.

| **Random forest** | **max depth** | **max features** | **n estimators** |
|---|---|---|---|
| **hi** | 10 | 10 | 40 |
| **tuned** | **182** | **311** | **319** |
| **lo** | 400 | 500 | 1200 |
| **Ada boosted decision tree** | **max depth** | **max features** | **n estimators** |
| **hi** | 10 | 10 | 10 |
| **tuned** | **442** | **476** | **221** |
| **lo** | 700 | 700 | 500 |
| **Multilayer Perceptron** | **hidden layer size** | **max iteration** | n/a |
| **hi** | 50 | 100 | n/a |
| **tuned** | **364** | **123** | |
| **lo** | 400 | 500 | |

Table S2. Raw Hemagglutination inhibition titers representing the homologous reference strain titer and heterologous test antigen titers.

| | A/swine/Indiana/A00968373/2012 | A/swine/Indiana/A00968373/2012 | A/swine/Iowa/A01480656/2014 | A/swine/Iowa/A01480656/2014 | A/swine/Pennsylvania/A01076777/2010 | A/swine/Pennsylvania/A01076777/2010 | A/swine/Indiana/A01202866/2011 | A/swine/Indiana/A01202866/2011 |
|---|---|---|---|---|---|---|---|---|
| A/swine/Indiana/A00968373/2012 | 640 | 2560 | | | | | | |
| A/swine/Nebraska/A01672826/2017 | 1280 | 2560 | | | | | | |
| A/swine/Iowa/A01480656/2014 | | | 1280 | 2560 | | | | |
| A/swine/Indiana/A02214844/2017 | | | 160 | 80 | | | | |
| A/swine/Pennsylvania/A01076777/2010 | | | | | 2560 | 640 | | |
| A/swine/North Carolina/A01732197/2016 | | | | | 320 | 160 | | |
| A/swine/Indiana/A01202866/2011 | | | | | | | 5120 | 5120 |
| A/swine/Iowa/A01733626/2016 | | | | | | | 40 | 80 |
| **Log2 Difference** | -1 | 0 | 3 | 5 | 3 | 2 | 7 | 6 |

Table S3. Random forest regression models feature importance for the 729 features used in the model. The importance of each predictor feature was calculated by the decrease in the node variance after fitting the random forest model. Feature position describes either pairwise HA1 domain percent identity, or location in the HA1 region, whereas Feature describes the specific positional mutation and its importance in the model.

| Feature position | Feature | importance |
|---|---|---|
| identity | identity | 0.31387702 |
| 145 | 145kn | 0.08126701 |
| 202 | 202iv | 0.05440468 |
| 222 | 222rw | 0.05151127 |
| 75 | 75hq | 0.02429311 |
| 137 | 137ry | 0.01509209 |
| 101 | 101dy | 0.01329186 |
| 62 | 62ek | 0.01045941 |
| 25 | 25il | 0.00992235 |
| 289 | 289ps | 0.00922882 |
| 133 | 133dn | 0.00770321 |
| 189 | 189ek | 0.00659735 |
| 92 | 92kt | 0.0064931 |
| 159 | 159hy | 0.00625947 |
| 50 | 50gr | 0.005632 |
| 121 | 121nt | 0.00546617 |
| 189 | 189kr | 0.00533521 |
| 133 | 133dg | 0.00517361 |
| 155 | 155hy | 0.00495734 |
| 160 | 160kr | 0.00424332 |
| 172 | 172de | 0.00386338 |
| 158 | 158kn | 0.00380091 |
| 189 | 189ks | 0.00375375 |
| 196 | 196av | 0.00372497 |
| 5 | 5eg | 0.00369966 |
| 193 | 193ns | 0.00369528 |
| 158 | 158gn | 0.00365699 |
| 227 | 227ps | 0.00361283 |
| 83 | 83ek | 0.00357943 |
| 138 | 138as | 0.00343168 |
| 196 | 196iv | 0.00337741 |
| 6 | 6ns | 0.003308 |
| 131 | 131ad | 0.00329187 |
| 144 | 144gv | 0.00320883 |
| 156 | 156kn | 0.00311055 |
| 140 | 140kr | 0.00309531 |
| 272 | 272av | 0.0030154 |
| 275 | 275dg | 0.00298937 |
| 242 | 242im | 0.00298038 |
| 261 | 261qr | 0.00295846 |
| 160 | 160kt | 0.00294396 |
| 175 | 175dn | 0.00293868 |
| 50 | 50er | 0.00287111 |
| 144 | 144nv | 0.0028249 |
| 186 | 186gs | 0.00272539 |
| 145 | 145ks | 0.00272259 |
| 156 | 156hs | 0.00264693 |
| 31 | 31dn | 0.00261551 |
| 220 | 220rv | 0.00259329 |
| 229 | 229ir | 0.00258695 |
| 111 | 111il | 0.00256987 |
| 46 | 46fs | 0.0025676 |
| 135 | 135st | 0.00255153 |
| 142 | 142eg | 0.0025191 |
| 203 | 203it | 0.00244917 |
| 217 | 217iv | 0.00240362 |
| 223 | 223iv | 0.00240271 |
| 276 | 276kn | 0.00232567 |
| 144 | 144dv | 0.0022111 |
| 273 | 273hp | 0.00220241 |
| 156 | 156hn | 0.00218761 |

| | | |
|---|---|---|
| 138 | 138at | 0.00218715 |
| 173 | 173kn | 0.00211613 |
| 50 | 50eg | 0.00206443 |
| 119 | 119ek | 0.00204617 |
| 48 | 48kt | 0.00204312 |
| 163 | 163av | 0.002004 |
| 246 | 246ns | 0.00194313 |
| 158 | 158en | 0.00191606 |
| 164 | 164lq | 0.00190432 |
| 163 | 163ae | 0.00189751 |
| 112 | 112iv | 0.0018937 |
| 122 | 122dn | 0.00188842 |
| 145 | 145ns | 0.00187564 |
| 46 | 46st | 0.00183688 |
| 158 | 158dn | 0.00182417 |
| 82 | 82kr | 0.00181536 |
| 38 | 38ns | 0.00181228 |
| 94 | 94fy | 0.00179581 |
| 262 | 262ns | 0.00175754 |
| 276 | 276en | 0.00174151 |
| 107 | 107st | 0.00173684 |
| 122 | 122nq | 0.00169302 |
| 159 | 159sy | 0.00168994 |
| 172 | 172dg | 0.00165302 |
| 190 | 190de | 0.00161775 |
| 142 | 142gk | 0.00161474 |
| 192 | 192it | 0.00160819 |
| 173 | 173kq | 0.00159163 |
| 299 | 299kr | 0.00159111 |
| 117 | 117nt | 0.00156774 |
| 142 | 142ek | 0.00156126 |
| 34 | 34iv | 0.00151198 |
| 273 | 273ps | 0.00150286 |
| 159 | 159fy | 0.00149139 |
| 156 | 156hk | 0.00149038 |
| 142 | 142gr | 0.00148088 |
| 137 | 137sy | 0.00146712 |
| 199 | 199ps | 0.00143094 |
| 140 | 140ik | 0.00143037 |
| 131 | 131at | 0.00142304 |
| 53 | 53dn | 0.00140467 |
| 262 | 262gs | 0.001402 |
| 193 | 193fs | 0.00139396 |
| 156 | 156ns | 0.00137745 |
| 118 | 118lm | 0.00136242 |
| 263 | 263eg | 0.00133284 |
| 3 | 3il | 0.00132069 |
| 189 | 189kn | 0.00129798 |
| 144 | 144iv | 0.00127793 |
| 155 | 155ny | 0.0012585 |
| 57 | 57qr | 0.00125616 |
| 185 | 185pt | 0.0012428 |
| 49 | 49gs | 0.0012341 |
| 45 | 45ns | 0.00122485 |
| 155 | 155ht | 0.00121085 |
| 53 | 53ns | 0.00119253 |
| 82 | 82ek | 0.00117609 |
| 194 | 194il | 0.00116949 |
| 311 | 311hq | 0.00116604 |
| 229 | 229iv | 0.00115827 |
| 165 | 165ns | 0.00115312 |
| 25 | 25ls | 0.00114176 |
| 197 | 197qr | 0.00112429 |
| 9 | 9ns | 0.0011199 |
| 226 | 226iv | 0.00110556 |
| 186 | 186iv | 0.00109807 |
| 157 | 157ls | 0.00109632 |
| 48 | 48at | 0.00108726 |
| 226 | 226il | 0.00108725 |
| 10 | 10mt | 0.00108277 |
| 138 | 138st | 0.0010668 |

| | | |
|---|---|---|
| 214 | 214is | 0.00105379 |
| 155 | 155ty | 0.00101471 |
| 106 | 106as | 0.00100503 |
| 309 | 309iv | 0.00099557 |
| 78 | 78dn | 0.00098015 |
| 124 | 124gs | 0.00097431 |
| 82 | 82kt | 0.00095214 |
| 182 | 182iv | 0.00094863 |
| 219 | 219fs | 0.00094022 |
| 214 | 214it | 0.00091132 |
| 233 | 233hy | 0.00090791 |
| 88 | 88iv | 0.00089785 |
| 262 | 262st | 0.00089779 |
| 137 | 137ny | 0.00088505 |
| 201 | 201ir | 0.0008789 |
| 133 | 133ns | 0.00087658 |
| 56 | 56hy | 0.0008492 |
| 150 | 150kr | 0.00084266 |
| 183 | 183hl | 0.00084256 |
| 78 | 78dg | 0.00082234 |
| 173 | 173kr | 0.00081493 |
| 82 | 82kn | 0.00080683 |
| 196 | 196tv | 0.00080468 |
| 212 | 212at | 0.00079831 |
| 124 | 124ds | 0.00079742 |
| 273 | 273hs | 0.00079663 |
| 229 | 229rv | 0.00079656 |
| 226 | 226iq | 0.00079052 |
| 135 | 135gt | 0.00078353 |
| 312 | 312kn | 0.00076105 |
| 2 | 2kn | 0.00075611 |
| 263 | 263gr | 0.00075153 |
| 186 | 186eg | 0.00075094 |
| 126 | 126dn | 0.00074237 |
| 307 | 307kr | 0.00073518 |
| 193 | 193sy | 0.00073224 |
| 189 | 189rs | 0.00072714 |
| 244 | 244fl | 0.00071948 |
| 189 | 189nr | 0.00071694 |
| 106 | 106av | 0.00070885 |
| 79 | 79fl | 0.00070488 |
| 203 | 203at | 0.00070407 |
| 149 | 149gs | 0.00070234 |
| 48 | 48it | 0.00069945 |
| 312 | 312ns | 0.00069408 |
| 58 | 58iv | 0.00069373 |
| 3 | 3lr | 0.00068643 |
| 260 | 260im | 0.00068315 |
| 7 | 7dg | 0.00068093 |
| 187 | 187st | 0.00067375 |
| 229 | 229gr | 0.00067085 |
| 228 | 228gs | 0.00065363 |
| 188 | 188dn | 0.00064955 |
| 189 | 189er | 0.00064895 |
| 328 | 328nt | 0.00064489 |
| 78 | 78eg | 0.00063968 |
| 207 | 207kr | 0.00063383 |
| 158 | 158ek | 0.00063061 |
| 263 | 263er | 0.00062696 |
| 275 | 275ds | 0.00062396 |
| 239 | 239ps | 0.00062292 |
| 11 | 11at | 0.0006225 |
| 208 | 208rw | 0.00062117 |
| 198 | 198as | 0.00061545 |
| 276 | 276ek | 0.00061035 |
| 225 | 225dg | 0.00060732 |
| 135 | 135as | 0.0006022 |
| 282 | 282it | 0.00059871 |
| 128 | 128nt | 0.00059603 |
| 198 | 198ae | 0.00058852 |
| 204 | 204iv | 0.00058677 |

| | | |
|---|---|---|
| 192 | 192at | 0.00058669 |
| 155 | 155hn | 0.00058199 |
| 137 | 137hy | 0.0005805 |
| 159 | 159qs | 0.00057959 |
| 167 | 167at | 0.00057379 |
| 137 | 137hs | 0.00057327 |
| 193 | 193rs | 0.00057113 |
| 78 | 78de | 0.0005687 |
| 156 | 156hq | 0.00055235 |
| 196 | 196ai | 0.00054881 |
| 144 | 144ns | 0.00053922 |
| 140 | 140ir | 0.00053855 |
| 146 | 146gs | 0.00052668 |
| 144 | 144de | 0.00052067 |
| 199 | 199is | 0.00052063 |
| 144 | 144av | 0.00051818 |
| 173 | 173nq | 0.0005179 |
| 128 | 128it | 0.00051405 |
| 45 | 45gs | 0.00051398 |
| 95 | 95ns | 0.00051189 |
| 133 | 133gn | 0.00051023 |
| 193 | 193ny | 0.00050941 |
| 131 | 131as | 0.00050836 |
| 225 | 225gn | 0.00050761 |
| 278 | 278kn | 0.00050326 |
| 156 | 156nq | 0.00050215 |
| 189 | 189ns | 0.0005015 |
| 124 | 124ns | 0.00049469 |
| 186 | 186gv | 0.00049446 |
| 248 | 248it | 0.00049143 |
| 144 | 144kv | 0.00049121 |
| 279 | 279fs | 0.00048954 |
| 213 | 213iv | 0.00048743 |
| 156 | 156ks | 0.00048634 |
| 159 | 159ny | 0.00048493 |
| 186 | 186ag | 0.00048383 |
| 144 | 144dn | 0.00048197 |
| 220 | 220rs | 0.00048167 |
| 186 | 186sv | 0.00047944 |
| 168 | 168mv | 0.00047889 |
| 201 | 201kr | 0.00047636 |
| 50 | 50kr | 0.00047373 |
| 142 | 142gs | 0.00046657 |
| 188 | 188de | 0.00046225 |
| 33 | 33qr | 0.00046158 |
| 193 | 193fn | 0.00045726 |
| 216 | 216ns | 0.00045416 |
| 82 | 82rt | 0.00044781 |
| 121 | 121it | 0.00044741 |
| 49 | 49dg | 0.00044584 |
| 135 | 135at | 0.00044083 |
| 144 | 144kn | 0.0004282 |
| 203 | 203ai | 0.0004281 |
| 144 | 144sv | 0.00042791 |
| 276 | 276nt | 0.00042244 |
| 278 | 278sy | 0.00041106 |
| 265 | 265gs | 0.00040904 |
| 133 | 133ds | 0.00040851 |
| 186 | 186is | 0.00040482 |
| 275 | 275gs | 0.00039562 |
| 128 | 128at | 0.00039559 |
| 105 | 105hy | 0.00039366 |
| 142 | 142er | 0.00039195 |
| 124 | 124dg | 0.00039149 |
| 1 | 1qr | 0.00039079 |
| 159 | 159fs | 0.00039065 |
| 53 | 53ds | 0.00039065 |
| 157 | 157lm | 0.00038988 |
| 117 | 117st | 0.00038843 |
| 3 | 3fl | 0.0003866 |
| 82 | 82ak | 0.0003854 |

| | | |
|---|---|---|
| 142 | 142kr | 0.00038385 |
| 278 | 278ns | 0.00037773 |
| 291 | 291dg | 0.00037706 |
| 227 | 227st | 0.00037525 |
| 144 | 144ev | 0.00037495 |
| 135 | 135gs | 0.00037134 |
| 278 | 278ny | 0.00036865 |
| 290 | 290hn | 0.00036495 |
| 328 | 328it | 0.00036333 |
| 262 | 262nt | 0.00036039 |
| 156 | 156kq | 0.00035956 |
| 21 | 21ps | 0.00035833 |
| 210 | 210hq | 0.00034761 |
| 225 | 225dn | 0.00034519 |
| 230 | 230iv | 0.00034393 |
| 208 | 208rs | 0.000337 |
| 172 | 172eg | 0.00033549 |
| 133 | 133de | 0.00032998 |
| 276 | 276kt | 0.00032849 |
| 192 | 192ai | 0.00032683 |
| 156 | 156ek | 0.000326 |
| 276 | 276ns | 0.00032423 |
| 112 | 112av | 0.00032388 |
| 244 | 244lv | 0.00032159 |
| 262 | 262gn | 0.00032145 |
| 94 | 94hy | 0.0003214 |
| 137 | 137cs | 0.00031437 |
| 209 | 209ns | 0.00031319 |
| 137 | 137rs | 0.00031231 |
| 143 | 143ps | 0.00030846 |
| 121 | 121in | 0.0003077 |
| 91 | 91ns | 0.00030443 |
| 83 | 83kn | 0.00030088 |
| 54 | 54ns | 0.00029825 |
| 158 | 158de | 0.00029723 |
| 29 | 29il | 0.00029714 |
| 124 | 124is | 0.0002938 |
| 159 | 159qy | 0.00029238 |
| 300 | 300iv | 0.00029093 |
| 144 | 144ad | 0.00028949 |
| 179 | 179iv | 0.00028774 |
| 199 | 199ip | 0.00028626 |
| 47 | 47st | 0.00028559 |
| 144 | 144in | 0.00028403 |
| 80 | 80kq | 0.00028393 |
| 196 | 196at | 0.00028179 |
| 83 | 83en | 0.00028049 |
| 210 | 210qr | 0.00027953 |
| 54 | 54rs | 0.00027942 |
| 144 | 144an | 0.00027927 |
| 326 | 326kr | 0.0002791 |
| 282 | 282iv | 0.00027858 |
| 304 | 304av | 0.00027823 |
| 173 | 173nr | 0.00027665 |
| 226 | 226qv | 0.00027489 |
| 114 | 114as | 0.00027362 |
| 165 | 165kn | 0.00027126 |
| 131 | 131an | 0.00027001 |
| 137 | 137cy | 0.00026894 |
| 271 | 271dn | 0.0002681 |
| 198 | 198at | 0.00026764 |
| 7 | 7de | 0.00026407 |
| 5 | 5gr | 0.0002636 |
| 78 | 78gn | 0.00026166 |
| 135 | 135et | 0.00025784 |
| 7 | 7ds | 0.00025591 |
| 193 | 193fy | 0.0002542 |
| 273 | 273hq | 0.0002541 |
| 215 | 215ps | 0.00025397 |
| 220 | 220kr | 0.00025077 |
| 51 | 51im | 0.00024991 |

| | | |
|---|---|---|
| 186 | 186as | 0.00024964 |
| 278 | 278dn | 0.00024947 |
| 287 | 287ns | 0.00024047 |
| 122 | 122dq | 0.00023824 |
| 159 | 159hs | 0.00023531 |
| 9 | 9rs | 0.00023154 |
| 160 | 160kq | 0.00023005 |
| 131 | 131ae | 0.00022965 |
| 81 | 81dn | 0.00022887 |
| 259 | 259kq | 0.00022694 |
| 159 | 159fq | 0.00022508 |
| 50 | 50ek | 0.00022252 |
| 264 | 264kr | 0.00021753 |
| 135 | 135ag | 0.00021651 |
| 172 | 172dn | 0.00021435 |
| 186 | 186gi | 0.00021412 |
| 144 | 144en | 0.00021292 |
| 323 | 323iv | 0.00020203 |
| 96 | 96dn | 0.00019959 |
| 6 | 6gn | 0.00019946 |
| 248 | 248nt | 0.00019897 |
| 193 | 193fr | 0.00019267 |
| 189 | 189es | 0.00018994 |
| 6 | 6gs | 0.00018902 |
| 142 | 142rs | 0.00018489 |
| 165 | 165en | 0.00018363 |
| 193 | 193kn | 0.00018066 |
| 201 | 201gr | 0.0001792 |
| 186 | 186av | 0.00017742 |
| 124 | 124di | 0.00017548 |
| 199 | 199as | 0.00017457 |
| 163 | 163at | 0.00017423 |
| 189 | 189kq | 0.00017188 |
| 167 | 167nt | 0.00017062 |
| 25 | 25fl | 0.00016849 |
| 131 | 131dt | 0.00016833 |
| 189 | 189gs | 0.00016447 |
| 93 | 93at | 0.00016433 |
| 6 | 6in | 0.0001611 |
| 144 | 144ae | 0.00016067 |
| 8 | 8dn | 0.00015799 |
| 31 | 31dg | 0.00015712 |
| 144 | 144dk | 0.00015663 |
| 156 | 156kr | 0.0001553 |
| 142 | 142gn | 0.00015517 |
| 310 | 310kr | 0.00015255 |
| 51 | 51il | 0.00015096 |
| 124 | 124dn | 0.00015081 |
| 197 | 197hq | 0.00014981 |
| 280 | 280ae | 0.00014906 |
| 238 | 238kr | 0.00014826 |
| 193 | 193fk | 0.00014763 |
| 144 | 144ik | 0.00014616 |
| 209 | 209gs | 0.00014284 |
| 48 | 48ai | 0.00014123 |
| 69 | 69as | 0.00014078 |
| 304 | 304at | 0.00014066 |
| 92 | 92kr | 0.00013856 |
| 264 | 264kn | 0.00013843 |
| 189 | 189gk | 0.00013415 |
| 224 | 224kr | 0.00013345 |
| 158 | 158dk | 0.00013329 |
| 135 | 135eg | 0.0001306 |
| 276 | 276et | 0.00013044 |
| 158 | 158gk | 0.00012903 |
| 156 | 156eh | 0.00012538 |
| 75 | 75hl | 0.00012472 |
| 82 | 82en | 0.00012368 |
| 276 | 276es | 0.00012265 |
| 248 | 248in | 0.00012192 |
| 144 | 144gn | 0.00011891 |

| | | |
|---|---|---|
| 117 | 117ns | 0.0001167 |
| 177 | 177lm | 0.00011651 |
| 173 | 173dk | 0.00011073 |
| 133 | 133gs | 0.00010982 |
| 278 | 278ks | 0.00010769 |
| 273 | 273pq | 0.00010768 |
| 193 | 193nr | 0.00010632 |
| 269 | 269kr | 0.0001055 |
| 137 | 137ns | 0.00010336 |
| 137 | 137nr | 0.0001004 |
| 50 | 50gk | 9.79E-05 |
| 196 | 196it | 9.74E-05 |
| 156 | 156qs | 9.70E-05 |
| 158 | 158dg | 9.63E-05 |
| 142 | 142ks | 9.41E-05 |
| 278 | 278in | 9.41E-05 |
| 246 | 246nt | 9.39E-05 |
| 226 | 226lv | 8.84E-05 |
| 144 | 144dg | 8.64E-05 |
| 159 | 159fn | 8.48E-05 |
| 124 | 124gn | 8.38E-05 |
| 156 | 156eq | 8.14E-05 |
| 189 | 189nq | 8.05E-05 |
| 304 | 304ad | 7.92E-05 |
| 131 | 131st | 7.81E-05 |
| 101 | 101df | 7.73E-05 |
| 216 | 216dn | 7.64E-05 |
| 31 | 31gn | 7.59E-05 |
| 3 | 3fi | 7.46E-05 |
| 121 | 121kn | 7.39E-05 |
| 214 | 214st | 7.33E-05 |
| 214 | 214iv | 7.31E-05 |
| 312 | 312ks | 7.26E-05 |
| 48 | 48ik | 7.20E-05 |
| 156 | 156en | 7.15E-05 |
| 131 | 131nt | 7.07E-05 |
| 144 | 144di | 6.86E-05 |
| 144 | 144gi | 6.86E-05 |
| 8 | 8kn | 6.68E-05 |
| 172 | 172en | 6.58E-05 |
| 275 | 275de | 6.56E-05 |
| 159 | 159hn | 6.47E-05 |
| 171 | 171kn | 6.47E-05 |
| 122 | 122qt | 6.33E-05 |
| 226 | 226lq | 6.26E-05 |
| 281 | 281cf | 6.26E-05 |
| 246 | 246st | 6.25E-05 |
| 128 | 128ai | 6.12E-05 |
| 208 | 208ks | 6.03E-05 |
| 78 | 78ds | 5.86E-05 |
| 62 | 62ei | 5.86E-05 |
| 124 | 124dr | 5.82E-05 |
| 172 | 172gn | 5.64E-05 |
| 329 | 329kr | 5.64E-05 |
| 292 | 292kr | 5.59E-05 |
| 247 | 247rs | 5.56E-05 |
| 20 | 20gv | 5.31E-05 |
| 208 | 208gs | 5.28E-05 |
| 57 | 57kr | 5.26E-05 |
| 250 | 250dn | 5.23E-05 |
| 144 | 144ks | 5.14E-05 |
| 160 | 160ak | 5.00E-05 |
| 144 | 144ag | 4.97E-05 |
| 106 | 106sv | 4.97E-05 |
| 124 | 124gi | 4.95E-05 |
| 203 | 203iv | 4.86E-05 |
| 276 | 276ks | 4.80E-05 |
| 137 | 137ch | 4.79E-05 |
| 83 | 83eg | 4.78E-05 |
| 62 | 62ik | 4.78E-05 |
| 169 | 169lp | 4.74E-05 |

| | | |
|---|---|---|
| 273 | 273lp | 4.73E-05 |
| 199 | 199ai | 4.72E-05 |
| 279 | 279ls | 4.72E-05 |
| 171 | 171dn | 4.71E-05 |
| 137 | 137hr | 4.55E-05 |
| 6 | 6is | 4.55E-05 |
| 229 | 229gi | 4.46E-05 |
| 133 | 133eg | 4.34E-05 |
| 159 | 159fh | 4.27E-05 |
| 144 | 144eg | 4.24E-05 |
| 207 | 207kq | 4.24E-05 |
| 273 | 273hl | 4.21E-05 |
| 208 | 208gr | 3.98E-05 |
| 135 | 135gk | 3.97E-05 |
| 83 | 83et | 3.97E-05 |
| 163 | 163ev | 3.86E-05 |
| 78 | 78en | 3.85E-05 |
| 131 | 131de | 3.83E-05 |
| 193 | 193as | 3.78E-05 |
| 173 | 173qr | 3.61E-05 |
| 203 | 203tv | 3.60E-05 |
| 163 | 163et | 3.55E-05 |
| 262 | 262gt | 3.55E-05 |
| 15 | 15lr | 3.52E-05 |
| 158 | 158eg | 3.51E-05 |
| 144 | 144ek | 3.41E-05 |
| 208 | 208kr | 3.21E-05 |
| 156 | 156hr | 3.13E-05 |
| 159 | 159ns | 3.10E-05 |
| 193 | 193ks | 3.10E-05 |
| 140 | 140kt | 3.07E-05 |
| 135 | 135kt | 2.99E-05 |
| 126 | 126nt | 2.95E-05 |
| 189 | 189en | 2.95E-05 |
| 83 | 83kt | 2.90E-05 |
| 227 | 227pt | 2.83E-05 |
| 159 | 159hq | 2.71E-05 |
| 189 | 189qr | 2.63E-05 |
| 122 | 122nt | 2.62E-05 |
| 21 | 21pq | 2.59E-05 |
| 275 | 275eg | 2.59E-05 |
| 101 | 101fy | 2.59E-05 |
| 2 | 2dk | 2.59E-05 |
| 124 | 124ir | 2.51E-05 |
| 193 | 193ry | 2.41E-05 |
| 25 | 25fi | 2.38E-05 |
| 327 | 327qr | 2.38E-05 |
| 62 | 62kr | 2.35E-05 |
| 220 | 220ir | 2.33E-05 |
| 189 | 189gn | 2.30E-05 |
| 83 | 83gk | 2.30E-05 |
| 144 | 144ds | 2.28E-05 |
| 135 | 135es | 2.15E-05 |
| 131 | 131ds | 2.15E-05 |
| 242 | 242iv | 2.15E-05 |
| 8 | 8ny | 2.09E-05 |
| 223 | 223av | 2.08E-05 |
| 122 | 122pq | 2.07E-05 |
| 2 | 2dn | 2.06E-05 |
| 142 | 142es | 2.05E-05 |
| 45 | 45gn | 1.99E-05 |
| 186 | 186ai | 1.98E-05 |
| 278 | 278ik | 1.97E-05 |
| 160 | 160ar | 1.97E-05 |
| 58 | 58il | 1.95E-05 |
| 6 | 6gr | 1.94E-05 |
| 214 | 214tv | 1.83E-05 |
| 94 | 94fh | 1.73E-05 |
| 57 | 57kq | 1.72E-05 |
| 197 | 197kq | 1.71E-05 |
| 208 | 208sw | 1.68E-05 |

| | | |
|---|---|---|
| 135 | 135ae | 1.64E-05 |
| 278 | 278is | 1.61E-05 |
| 208 | 208ir | 1.58E-05 |
| 278 | 278dy | 1.58E-05 |
| 193 | 193an | 1.54E-05 |
| 124 | 124gr | 1.53E-05 |
| 48 | 48ak | 1.52E-05 |
| 25 | 25is | 1.48E-05 |
| 167 | 167tv | 1.46E-05 |
| 131 | 131en | 1.46E-05 |
| 78 | 78gs | 1.41E-05 |
| 133 | 133en | 1.36E-05 |
| 199 | 199ap | 1.23E-05 |
| 173 | 173dq | 1.19E-05 |
| 6 | 6rs | 1.16E-05 |
| 142 | 142kn | 1.13E-05 |
| 137 | 137fy | 1.12E-05 |
| 121 | 121kt | 1.11E-05 |
| 131 | 131ns | 1.06E-05 |
| 156 | 156nr | 1.06E-05 |
| 189 | 189gr | 1.04E-05 |
| 160 | 160at | 1.04E-05 |
| 32 | 32de | 1.01E-05 |
| 260 | 260lm | 9.75E-06 |
| 124 | 124rs | 9.30E-06 |
| 144 | 144gk | 9.17E-06 |
| 62 | 62er | 9.17E-06 |
| 302 | 302fy | 9.12E-06 |
| 9 | 9nr | 8.62E-06 |
| 92 | 92rt | 8.53E-06 |
| 198 | 198es | 8.52E-06 |
| 6 | 6nr | 8.44E-06 |
| 3 | 3ir | 8.07E-06 |
| 273 | 273qs | 8.04E-06 |
| 140 | 140rt | 7.98E-06 |
| 189 | 189qs | 7.96E-06 |
| 156 | 156er | 7.81E-06 |
| 193 | 193ay | 7.31E-06 |
| 261 | 261kq | 7.26E-06 |
| 197 | 197kr | 7.09E-06 |
| 260 | 260il | 6.69E-06 |
| 82 | 82at | 6.49E-06 |
| 278 | 278dk | 6.30E-06 |
| 156 | 156rs | 6.01E-06 |
| 75 | 75lq | 5.95E-06 |
| 186 | 186es | 5.70E-06 |
| 144 | 144ai | 5.52E-06 |
| 142 | 142nr | 5.47E-06 |
| 45 | 45is | 5.23E-06 |
| 126 | 126dt | 5.23E-06 |
| 261 | 261kr | 5.10E-06 |
| 142 | 142en | 5.02E-06 |
| 156 | 156es | 4.89E-06 |
| 137 | 137hn | 4.86E-06 |
| 62 | 62ir | 4.56E-06 |
| 144 | 144ei | 4.52E-06 |
| 137 | 137fn | 4.25E-06 |
| 160 | 160qr | 4.04E-06 |
| 278 | 278ky | 4.02E-06 |
| 122 | 122np | 3.85E-06 |
| 144 | 144ak | 3.78E-06 |
| 63 | 63dn | 3.50E-06 |
| 137 | 137cn | 3.48E-06 |
| 223 | 223ai | 2.84E-06 |
| 137 | 137cr | 2.66E-06 |
| 278 | 278ds | 2.51E-06 |
| 173 | 173dn | 2.20E-06 |
| 160 | 160rt | 1.97E-06 |
| 193 | 193kr | 1.89E-06 |
| 198 | 198st | 1.37E-06 |
| 193 | 193af | 1.34E-06 |

| | | |
|---|---|---|
| 135 | 135ak | 1.30E-06 |
| 156 | 156qr | 1.29E-06 |
| 208 | 208is | 1.26E-06 |
| 135 | 135ks | 1.15E-06 |
| 159 | 159nq | 9.90E-07 |
| 207 | 207qr | 8.91E-07 |
| 137 | 137fs | 7.26E-07 |
| 186 | 186ev | 1.81E-07 |
| 45 | 45in | 3.39E-08 |
| 259 | 259qr | 0 |
| 220 | 220sv | 0 |
| 8 | 8dk | 0 |
| 46 | 46ft | 0 |
| 137 | 137cf | 0 |
| 264 | 264nr | 0 |
| 259 | 259kr | 0 |
| 82 | 82nr | 0 |
| 282 | 282tv | 0 |
| 275 | 275es | 0 |
| 216 | 216ds | 0 |
| 49 | 49ds | 0 |
| 140 | 140it | 0 |
| 82 | 82er | 0 |
| 82 | 82et | 0 |
| 131 | 131et | 0 |
| 214 | 214sv | 0 |
| 160 | 160aq | 0 |
| 220 | 220ks | 0 |
| 122 | 122dt | 0 |
| 122 | 122dp | 0 |
| 193 | 193ak | 0 |
| 244 | 244fv | 0 |
| 82 | 82nt | 0 |
| 167 | 167av | 0 |
| 279 | 279fl | 0 |
| 6 | 6ir | 0 |
| 124 | 124in | 0 |
| 208 | 208gw | 0 |
| 6 | 6gi | 0 |
| 83 | 83gn | 0 |
| 131 | 131es | 0 |
| 78 | 78es | 0 |
| 121 | 121ik | 0 |
| 144 | 144gs | 0 |
| 160 | 160qt | 0 |
| 189 | 189gq | 0 |
| 45 | 45gi | 0 |
| 137 | 137fr | 0 |
| 112 | 112ai | 0 |
| 3 | 3fr | 0 |
| 278 | 278di | 0 |
| 208 | 208gi | 0 |
| 163 | 163tv | 0 |
| 186 | 186ei | 0 |
| 273 | 273lq | 0 |
| 122 | 122pt | 0 |
| 189 | 189eq | 0 |
| 201 | 201gk | 0 |
| 83 | 83gt | 0 |
| 8 | 8ky | 0 |
| 186 | 186ae | 0 |
| 188 | 188en | 0 |
| 131 | 131dn | 0 |
| 58 | 58lv | 0 |
| 128 | 128in | 0 |
| 165 | 165ek | 0 |
| 155 | 155nt | 0 |
| 193 | 193ar | 0 |
| 142 | 142ns | 0 |
| 82 | 82ae | 0 |
| 144 | 144as | 0 |

| | | |
|---|---|---|
| 144 | 144es | 0 |
| 137 | 137fh | 0 |
| 189 | 189eg | 0 |
| 203 | 203av | 0 |
| 157 | 157ms | 0 |
| 278 | 278iy | 0 |
| 135 | 135ek | 0 |
| 54 | 54nr | 0 |
| 133 | 133es | 0 |
| 21 | 21qs | 0 |
| 128 | 128an | 0 |
| 82 | 82ar | 0 |
| 276 | 276st | 0 |
| 229 | 229gv | 0 |
| 208 | 208gk | 0 |
| 144 | 144is | 0 |
| 165 | 165es | 0 |
| 210 | 210hr | 0 |
| 124 | 124nr | 0 |
| 198 | 198et | 0 |
| 5 | 5er | 0 |
| 197 | 197hr | 0 |
| 201 | 201ik | 0 |
| 220 | 220is | 0 |
| 273 | 273ls | 0 |
| 193 | 193ky | 0 |
| 167 | 167an | 0 |
| 83 | 83nt | 0 |
